# Annotation of biologically relevant ligands in UniProtKB using ChEBI

**DOI:** 10.1101/2022.08.19.504519

**Authors:** Elisabeth Coudert, Sebastien Gehant, Edouard de Castro, Monica Pozzato, Delphine Baratin, Teresa Batista Neto, Christian J.A. Sigrist, Nicole Redaschi, Alan Bridge, The UniProt Consortium

## Abstract

**Motivation:** To provide high quality, computationally tractable annotation of binding sites for biologically relevant (cognate) ligands in UniProtKB using the chemical ontology ChEBI (Chemical Entities of Biological Interest), to better support efforts to study and predict functionally relevant interactions between proteins and small molecule ligands.

**Results:** We structured the data model for cognate ligand binding site annotations in UniProtKB and performed a complete reannotation of all cognate ligand binding sites using stable unique identifiers from ChEBI, which we now use as the reference vocabulary for all such annotations. We developed improved search and query facilities for cognate ligands in the UniProt website, REST API and SPARQL endpoint that leverage the chemical structure data, nomenclature, and classification that ChEBI provides.

**Availability:** Binding site annotations for cognate ligands described using ChEBI are available for UniProtKB protein sequence records in several formats (text, XML, and RDF), and are freely available to query and download through the UniProt website (www.uniprot.org), REST API (www.uniprot.org/help/api), SPARQL endpoint (sparql.uniprot.org/), and FTP site (https://ftp.uniprot.org/pub/databases/uniprot/).

**Supplementary information:** Supplementary Table 1.

## 1 Introduction

The UniProt Knowledgebase (UniProtKB, at www.uniprot.org) is a reference resource of protein sequences and functional annotation that covers proteins from all branches of the tree of life (The UniProt Consortium, 2021). UniProtKB includes an expert curated core of around 567,000 reviewed UniProtKB/Swiss-Prot protein sequence entries and over 226 million unreviewed UniProtKB/TrEMBL entries that are annotated by automatic systems (MacDougall, et al., 2020) (statistics for release 2022_03 of August 2022). UniProtKB provides a wealth of information on protein sequences and their functions, including descriptions of the nature and binding sites of biologically relevant or “cognate” ligands (the term used in the remainder of this paper) (Das and Orengo, 2018; Tyzack, et al., 2018) such as activators, inhibitors, cofactors, and substrates. UniProt curators capture knowledge of cognate ligands through expert literature curation and from experimentally resolved protein structures in the Protein Data Bank (PDB/PDBe) (Armstrong, et al., 2020; Burley, et al., 2021; Velankar, et al., 2021), removing adventitious ligands that are technical artefacts and mapping experimentally observed ligands to their cognate equivalents when necessary.

In this work, we describe improvements to the annotation of cognate ligands and their binding sites in UniProtKB using the chemical ontology ChEBI (Chemical Entities of Biological Interest, www.ebi.ac.uk/chebi/) (Hastings, et al., 2016). We have performed a complete reannotation of all cognate ligand binding sites in UniProtKB, replacing textual descriptions of defined ligands with stable unique identifiers from the ChEBI ontology, and now use ChEBI as the reference vocabulary for all new ligand binding site annotations. This new dataset will improve interoperability of UniProtKB with other resources of cognate ligand binding site annotations such as PDBe (Mukhopadhyay, et al., 2019), BioLiP (Yang, et al., 2013), FireDB (Maietta, et al., 2014), MetalPDB (Putignano, et al., 2018), and PDBBind (Liu, et al., 2015), and provide better support for efforts to study, and predict, functional interactions between proteins and their cognate ligands using computational approaches (Das, et al., 2021; Littmann, et al., 2021; Wehrspan, et al., 2021; Wu, et al., 2018).

## 2 Methods

### 2.1 Changes to the UniProt data model and formats

Most sequence annotations (also called “features”) in UniProtKB, including cognate ligand binding site annotations that are the subject of this work, consist of three main elements. The “feature location” defines the sequence region or amino acid residue position that is annotated, the “feature key” specifies the type of each feature, and the “feature description” provides a textual description, which for cognate ligands includes the name of the ligand and other relevant information, such as numbering (of multiple ligands of the same type) and ligand roles. We structured this description for binding site annotations into several fields (described in the online documentation at www.uniprot.org/release-notes/2022-08-03-release), to standardize the description of a ligand, and optionally the bound part of the ligand (such as the iron atom in a heme, or a nucleotide in a macromolecule such as DNA), with the ChEBI ontology. We illustrate this new data model with examples in the Results section. We also simplified the range of feature keys that are used for cognate ligand binding site annotations, which were the following:

- “CA_BIND”, which denotes a sequence region that binds to calcium;
- “METAL”, which denotes a sequence position that binds a metal;
- “NP_BIND”, which denotes a sequence region that binds a nucleotide phosphate;
- “BINDING”, which denotes a sequence position that binds any type of chemical entity;
- “REGION”, which denotes a sequence region of interest in a protein (including a region that binds a ligand).

The ChEBI ontology provides a means to search for any ligand or class of ligand represented in ChEBI, at any desired level of specificity, without requiring ligand-specific feature keys. We therefore deprecated the feature keys “CA_BIND”, “METAL”, and “NP_BIND”, and now use the feature key “BINDING” for all binding site annotations for all cognate ligands. We also recurated all cognate ligand binding sites of interest found in features of the type “REGION”, and moved them to “BINDING”. Finally, we modified all UniRules (MacDougall, et al., 2020), including HAMAP (Pedruzzi, et al., 2015) and PROSITE (Sigrist, et al., 2013) rules, to provide binding site annotations using ChEBI identifiers in the new data model described here.

### 2.2 Mapping of legacy text annotations of cognate ligand binding sites in UniProtKB to ChEBI

UniProtKB previously described cognate ligands in binding site annotations using text labels, such as ‘lipid’, ‘cholesterol’, ‘heme’, ‘heme b’, ‘divalent metal’, or ‘zinc’. To standardize the descriptions of biologically relevant ligands in binding site annotations in UniProtKB we created a one-to-one mapping between each such text label and the corresponding ChEBI identifier, and used that mapping to reannotate all legacy data.

We extracted unique ligand descriptions from binding site annotations linked to each of the feature keys “CA_BIND”, “METAL”, “NP_BIND”, and “BINDING”, as well as “REGION” annotations with the word “binding” in the feature description, and mapped each of the text labels found to the corresponding ChEBI identifier manually. During the mapping we selected the ChEBI that represents the major microspecies of the ligand (the predominant protonation state) at pH7.3, which is the convention used in UniProtKB and the Rhea reaction knowledgebase (www.rhea-db.org) (Bansal, et al., 2022). Some ligand text labels presented with multiple possible mappings to ChEBI – sometimes due to stereochemistry issues – while some described generic classes or roles such as cofactor, hormone, or odorant, which we could not map to any defined structure. We examined each of these cases in turn and, where necessary, recurated them, using information from the literature, the PDB and the UniProtKB protein sequence records concerned, including existing Rhea reaction annotations, before selecting the most appropriate mapping to ChEBI.

Once complete, we used the mapping of defined cognate ligands to replace legacy text labels in UniProtKB with the corresponding identifiers from ChEBI. We also used additional information from the existing annotations, such as ligand numbering and roles, to populate the corresponding data fields in the new structured data model.

We did not yet systematically recurate binding site annotations for enzymes in UniProtKB with the generic text label “substrate”, which does not specify which of the possible substrate(s) are bound. We are continuing to map these legacy “substrate” annotations to specific ChEBI identifiers, using Rhea annotations and other information such as ligand data from PDBe records where available, mapped to UniProt sequences using the SIFTS framework (Dana, et al., 2019).

### 2.3 UniProt tools and services to exploit ligand binding site annotations

We modified the UniProt website www.uniprot.org, UniProt REST API www.uniprot.org/help/api, and UniProt SPARQL endpoint sparql.uniprot.org/, to support searches for ligand binding site annotations using ChEBI identifiers, ligand names, synonyms, and chemical structures from ChEBI encoded as InChIKeys. The InChIKey is a simple hash representation of chemical structures that provides a convenient means to search and map chemical structure databases (see www.inchi-trust.org/).

## 3 Results

### 3.1 Structuring cognate ligand binding site annotations in UniProtKB using ChEBI

The annotation of cognate ligand binding sites in UniProtKB using the chemical ontology ChEBI was made available from UniProt release 2022_03 of August 2022. It features 776 unique ligands from ChEBI, which are involved in over 980,000 binding site annotations for over 200,000 UniProtKB/Swiss-Prot protein sequence records, and over 65 million binding site annotations for over 17 million protein sequence records for the whole of UniProtKB, including UniProtKB/TrEMBL. We provide a complete list of all cognate ligands used in UniProtKB release 2022_03 in Supplementary Table 1.

The new data model improves consistency of annotations while retaining flexibility. It allows annotations at any level of granularity in ChEBI, from broad classes of ligand such as “metal cation” (CHEBI:25213) or “heme” (CHEBI:30413) to structurally defined ligands such as “Fe(2+)” (CHEBI:29033) or “heme b” (CHEBI:60344). It supports annotation of binding sites for small molecules, which constitute the vast majority of ligands curated in UniProtKB, and also for parts of larger macromolecules, such as the iron atom (CHEBI:18248) bound as part of a heme b (CHEBI:60344) as in the example below, taken from UniProtKB/Swiss-Prot entry P00175 in text format. This example also shows the “evidence” field which lists the evidences that support the annotation, each described by a term from the Evidence and Conclusions Ontology ECO (Nadendla, et al., 2022), and the source of the information, here experiments published in two peer reviewed articles (Cunane, et al., 2002; Xia and Mathews, 1990) and protein structures 1FCB and 1KBI from the PDB.

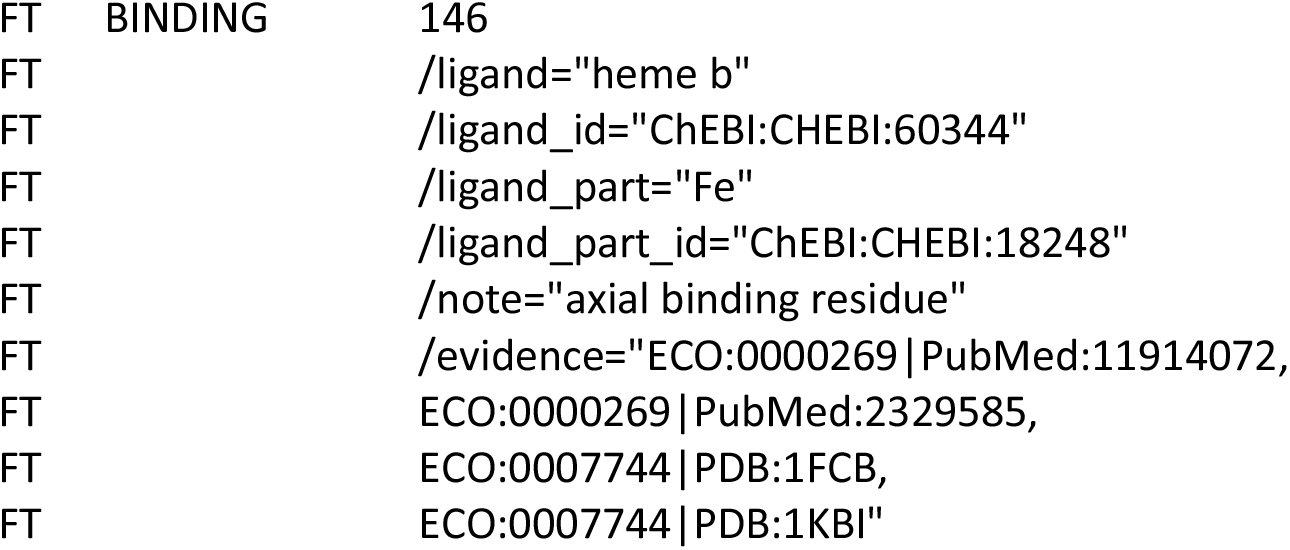

We refer readers to the online documentation at www.uniprot.org/release-notes/2022-08-03-release, which provides additional examples of binding site annotations in the UniProtKB formats text, XML, and RDF/XML.

### 3.2 UniProt tools and services to access and query cognate ligand binding site annotations made with ChEBI

Users can access and query UniProtKB ligand binding site annotations made with ChEBI using the UniProt website, REST API and SPARQL endpoint.

#### 3.2.1 UniProt website

The UniProt website www.uniprot.org provides access to UniProtKB protein sequence records and annotations, including cognate ligand binding site annotations for each protein (Figure 1). Users can now query the website for proteins that bind ligands of interest using identifiers, names, synonyms, and chemical structures (encoded as InChIKeys) from ChEBI using the advanced query builder. The complete ChEBI ontology is indexed, so that searches using identifiers for higher level grouping classes in the ChEBI ontology will retrieve UniProt records with binding site annotations to all child classes. ChEBI identifiers entered by users are automatically mapped to those of the major microspecies at pH 7.3, which is the form used in UniProtKB and Rhea, using a mapping file provided by Rhea.

**Fig. 1.**
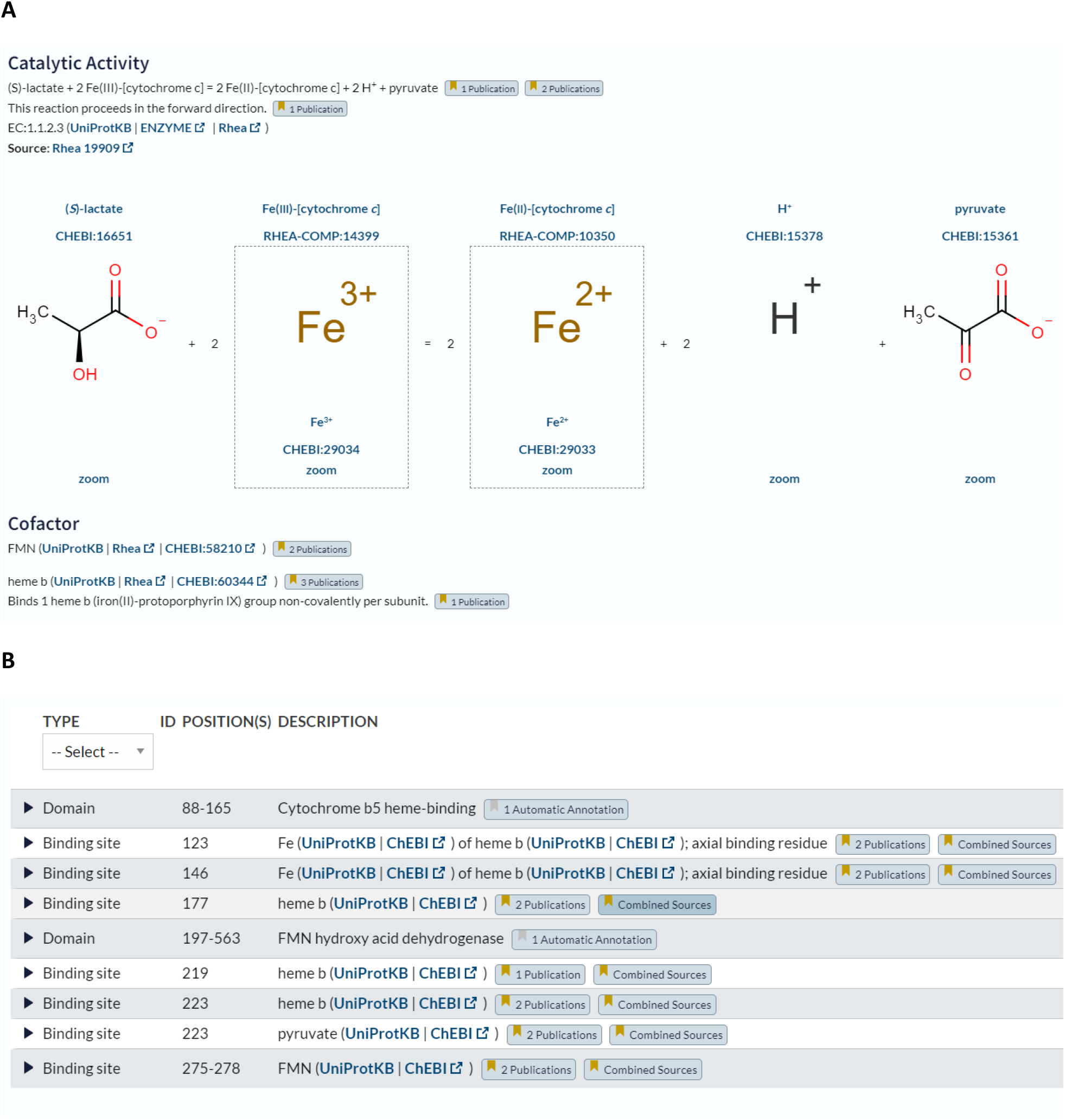
Website view of small molecule annotations in UniProtKB – from www.uniprot.org/uniprotkb/P00175/entry. All small molecule annotations are shown in the “Function” section. **(A)** The “Catalytic Activity” subsection describes enzymatic reactions using Rhea (which is based on ChEBI), while cofactors are described using ChEBI in the “Cofactor” subsection. Standardization of reaction and cofactor descriptions was performed in previous work, and is shown here for completeness. (**B**) The “Features” subsection displays the available binding site annotations for cognate ligands described using ChEBI, the subject of this work. Each ligand has a link “UniProtKB” to launch searches for other proteins binding this ligand, a link out to “ChEBI”, and expandable sections like “Publications” to examine provenance and evidence.

The sample query shown below can be typed into the search box: it will retrieve all proteins with binding site annotations for any kind of heme, using the ChEBI identifier for that grouping class, which is ChEBI:30413:

1. (ft_binding:”CHEBI:30413”) The result set will include all proteins with binding site annotations for any form of heme (CHEBI:30413), including “heme b” (CHEBI:60344), “heme c” (CHEBI:61717), and all others. To retrieve proteins with binding site annotations for specific forms of heme, users can simply change the ChEBI identifier to that of the form desired, here ChEBI:60344 for “heme b”:
2. (ft_binding:”CHEBI:60344”) Users can also perform searches for specific ligands using the chemical structure represented as an InChIKey, if using a chemical structure database other than ChEBI:
3. (ft_binding:KABFMIBPWCXCRK-RGGAHWMASA-J) Users can elect to ignore charge, by removing the third block of the InChIKey, as in this example:
4. (ft_binding:KABFMIBPWCXCRK-RGGAHWMASA) They may also elect to ignore both stereochemistry and charge, by removing both the second and third blocks of the InChIKey:
5. (ft_binding:KABFMIBPWCXCRK) Users can also combine searches for ligand binding site annotations with other types of annotations, as in this query for human mitochondrial flavoproteins (i.e. proteins with annotated binding sites for some CHEBI:30527 - flavin) that are linked to genetic diseases defined by the resource OMIM (Online Mendelian Inheritance in Man) (Hamosh, et al., 2021):
6. (ft_binding:”CHEBI:30527”) AND (cc_scl_term:SL-0173) AND (organism_id:9606) AND (cc_disease:*)

We provide complete documentation on searching for small molecule data in UniProtKB, including ligands described in binding site annotations, at www.uniprot.org/help/chemical_data_search.

#### 3.2.2 UniProt REST API

The UniProt REST API (www.uniprot.org/help/api) allows users to query and process UniProt data programmatically and to specify the required output format for query results (such as txt, xml, rdf, tsv, etc.) and, for tab-separated format, the desired annotation fields. The simplest way to create URLs for programmatic use is by using the advanced query builder to set the desired query fields and values, perform the search and click the “Download” button, which opens a panel with a “Generate URL for API” link. Users can now query the UniProt REST API with identifiers, names, synonyms and chemical structures from ChEBI for ligand binding site annotations.

#### 3.2.3 UniProt SPARQL endpoint

The UniProt SPARQL endpoint sparql.uniprot.org allows users to query UniProt RDF data and RDF data from other SPARQL endpoints using federated SPARQL queries. It now supports queries for ligand binding site annotations using identifiers, names, synonyms and chemical structure data from ChEBI. We demonstrate this capability using a federated SPARQL query that combines the UniProt SPARQL endpoint and that of the Integrated Database of Small Molecules (IDSM) (Galgonek and Vondrasek, 2021; Kratochvil, et al., 2019). IDSM supports fingerprint-guided chemical similarity and substructure searches in a number of chemical datasets, including ChEBI, using Sachem, a high performance open source chemical cartridge (Kratochvil, et al., 2018). This federated SPARQL query allows UniProt to borrow that functionality from IDSM; it will find all proteins that bind to ligands with structures similar to that of a query ligand, in this case heme b (specified using SMILES, or Simplified Molecular-Input Line-Entry notation) (http://opensmiles.org). The UniProt SPARQL endpoint queries that of IDSM, which returns the set of chemical entities in ChEBI that are similar to the query ligand heme b, and then searches for proteins in UniProtKB with binding site annotations for those ligands, which it then returns to the user.

~~~
SELECT ?uniprot ?mnemonic ?proteinName ?ligandSimilarityScore ?ligand
WHERE {
  SERVICE <https://idsm.elixir-czech.cz/sparql/endpoint/chebi> {
   [sachem:compound ?ligand; sachem:score ?ligandSimilarityScore]
   sachem:similaritySearch
   [
   sachem:query
    “CC1=C(CCC([O-])=O)C2=[N+]3C1=Cc1c(C)c(C=C)c4C=C5C(C)=C(C=C)C6=[N+]5[Fe-
    ]3(n14)n1c(=C6)c(C)c(CCC([O-])=O)c1=C2”;
    sachem:cutoff “8e-1”^^xsd:double ;
    sachem:aromaticityMode sachem:aromaticityDetect ;
    sachem:similarityRadius 1 ;
    sachem:tautomerMode sachem:ignoreTautomers
  ]
}
?uniprot up:mnemonic ?mnemonic .
?uniprot up:recommendedName/up:fullName ?proteinName .
?uniprot up:annotation ?annotation .
?annotation a up:Binding_Site_Annotation ;
up:ligand/rdfs:subClassOf ?ligand .
}
ORDER BY DESC(?ligandSimilarityScore)
~~~

We provide more sample queries in the online documentation for the UniProt SPARQL endpoint at https://sparql.uniprot.org/.well-known/sparql-examples/. Additional documentation on the IDSM SPARQL endpoint is provided at https://idsm.elixir-czech.cz/sparql/doc/manual.html.

## 4 Conclusions and future work

We have structured and reannotated cognate ligand binding sites in UniProtKB using ChEBI, which is now the standard vocabulary for all small molecule ligand annotations in UniProtKB, and report new tools and services to exploit this improved ligand dataset via the UniProt website and APIs. This work is part of an ongoing program to standardize all small molecule annotations in UniProtKB using ChEBI, and builds on previous improvements to enzyme and transporter annotation in UniProtKB using the Rhea knowledgebase of biochemical reactions, which uses ChEBI to represent reactants (Bansal, et al., 2022; Morgat, et al., 2020). We are continuing to work to improve the UniProt ligand dataset through expert curation of cognate ligands and their binding sites from the literature, and are currently working to develop improved pipelines for import and curation of ligand data from PDBe using curated mappings of experimental ligands in PDBe to cognate ligands in ChEBI.

## Acknowledgements

We thank the Cheminformatics and Metabolism Team of EMBL-EBI for their work in maintaining and developing ChEBI, particularly Adnan Malik and Gareth Owen for expert curation and advice and Andrew Leach, and the PDBe team at EMBL-EBI, for maintaining and developing PDBe, the principal source of ligand data for UniProtKB. We also thank Sameer Velankar of PDBe at EMBL-EBI for stimulating discussions on methods to identify cognate ligands in protein structures in PDBe. We gratefully acknowledge the software contributions of ChemAxon (https://www.chemaxon.com/products/marvin/).

